# Col-Ovo: Smartphone-based artificial intelligence for rapid counting of *Aedes* mosquito eggs under field conditions

**DOI:** 10.64898/2026.03.19.712860

**Authors:** Juan Almanza, Diego Montenegro

## Abstract

**Background:** OviCol has recently been proposed as a disruptive strategy for the surveillance and control of synanthropic *Aedes* mosquitoes, vectors of dengue, Zika, and chikungunya viruses. The approach integrates monitoring and control through ultra–low-cost ovitraps (∼0.2 USD), bioattractants, and egg inactivation using hot water. However, large-scale ovitrap surveillance generates thousands of egg substrates that require time-consuming manual counting, creating a major operational bottleneck. To address this limitation, we developed Col-Ovo, an artificial intelligence–based tool for automated counting of *Aedes aegypti* eggs from real field samples, together with OviLab, a digital platform for annotation, curation, and management of entomological image datasets.

**Methodology/Principal Findings:** The detection model was trained using YOLOv11m on a dataset of 275 oviposition substrates (20.5 cm strips) collected under routine operational conditions. Images were captured in situ without preprocessing and included substrates heavily stained by bioattractants such as blackstrap molasses and dry yeast (*Saccharomyces cerevisiae*), as well as sand and particulate debris, reflecting realistic field conditions. The system was designed to operate with standard smartphone images and tolerate compression artifacts produced by messaging platforms such as WhatsApp. Performance was evaluated by comparing automated egg counts with expert manual counts and with virtual-human counts conducted in OviLab using >200% image magnification. Col-Ovo achieved >95% agreement with expert counts and 88% agreement with OviLab while reducing processing time from approximately 15 minutes to <3 seconds per sample.

**Conclusions/Significance:** Col-Ovo enables rapid, scalable quantification of *Ae. aegypti* eggs from smartphone images, addressing a critical operational barrier in ovitrap-based surveillance. The system requires no image preprocessing or specialized hardware and is accessible through a lightweight web interface supported by an AI architecture that allows retraining for new ecological contexts or additional *Aedes* species. Integrated with OviLab, this platform provides a flexible digital infrastructure that can strengthen routine vector surveillance and community-level control programs across regions where *Aedes* mosquitoes continue to expand.

**Author Summary:** Mosquitoes that transmit dengue, Zika, and chikungunya are expanding in many parts of the world. Monitoring their populations is essential for guiding prevention and control actions. A common surveillance method uses small traps where female mosquitoes lay their eggs. By counting the eggs collected in these traps, health programs can estimate mosquito abundance and detect increases in risk. However, the eggs are extremely small: about 0.065 mm², and are usually counted manually under magnification. This process is slow, requires trained personnel, and limits how many samples can be analyzed in routine surveillance.

In this study, we developed a digital tool that automatically counts mosquito eggs from photographs taken with a smartphone. The system was trained using images collected under real field conditions, including samples with stains, dirt, and other materials commonly found in mosquito traps. The tool can analyze images even when they are compressed and shared through WhatsApp. By reducing counting time from 15 minutes to only a 25 seconds, this approach can help strengthen mosquito surveillance and support faster responses to mosquito-borne disease risks.

## Introduction

Mosquitoes of the family *Culicidae* are the deadliest animals worldwide, accounting for more than 725,000 deaths annually [1]. Among them, synanthropic species such as *Aedes aegypti* and *Aedes albopictus* play a major role in the transmission of arboviruses of global public health importance, including dengue, chikungunya, Zika, and urban yellow fever [2,3]. In recent years, the re-emergence of urban yellow fever in several countries in the Americas has generated additional public health concern, with reported case fatality rates reaching 40.5% (53/131) in 2025 [4].

Over the past decades, these mosquitoes have expanded far beyond their historical ranges. Today, both *Aedes* species are established in approximately 170 countries worldwide[5,6], reflecting their remarkable ecological adaptability and the growing interconnectedness of global trade and urbanization. This expansion has been accompanied by a substantial economic burden. Recent estimates indicate that the global costs associated with *Aedes*-borne diseases fluctuate with epidemic cycles, ranging between 3.1 and 20.3 trillion USD annually when accounting for direct healthcare expenses, productivity losses, and long-term disability [7].

Together, these trends underscore the urgent need to strengthen and diversify vector surveillance, prevention, and control strategies, particularly in resource-limited settings where arboviral transmission is most intense.

Within this context, ovitrap-based surveillance has emerged as an important tool for monitoring *Aedes* populations and detecting early vector activity [8]. More recently, low-cost handmade ovitraps incorporating bioattractants and hot water treatment have been proposed as a cost-effective community-based strategy for eliminating *Aedes* eggs [9]. This approach (known as OviCol) is based on the premise that targeting immature stages of the vector can be more cost-efficient, environmentally sustainable, and operationally feasible than relying exclusively on adulticide interventions. The incorporation of OviCol into routine vector surveillance programs in several endemic regions of Colombia has demonstrated its operational feasibility and effectiveness, achieving results comparable—and in some cases superior—to those obtained using conventional mass-produced ovitraps [10].

Despite these advantages, a key limitation remains the rapid and reliable quantification of eggs deposited on oviposition substrates. Manual counting by trained personnel remains the standard approach but it is labor-intensive, susceptible to human error, and difficult to scale across large surveillance networks. This constraint limits the potential of ovitrap-based strategies as continuous and community-level surveillance tools.

Artificial intelligence (AI) has therefore been proposed as a potential solution to automate egg detection and counting. Several systems have been developed; however, most remain confined to research environments rather than routine vector control settings. For example, MecVision [11] showed limited agreement (17.0%) with expert human counts under field conditions [9].

EggCountAI [12], based on convolutional neural networks, reported high accuracy under laboratory conditions but requires preprocessed images, computer-based operation, and relatively complex interfaces, reducing its suitability for community-level implementation. Similarly, Ovitrap Monitor [13] proposed compatibility with smartphone images; however, the system was not publicly accessible at the time of this study, limiting reproducibility and operational adoption.

More broadly, most AI-based models for *Aedes* egg recognition have been trained using laboratory datasets under controlled imaging conditions. Their performance in real-world environments—characterized by contaminated substrates (dust, sand, organic stains, or non-target insect eggs), heterogeneous materials, and variable lighting—remains uncertain. Additional constraints include restriction to narrow wooden paddles (<3 cm wide), dependence on external image preprocessing software, and limited capacity to analyze photographs captured directly in situ using mobile phones [14–16]. As a result, many of these systems function effectively as research prototypes but remain poorly adapted to the operational realities of routine surveillance and vector control programs.

To address these limitations, we developed an IA system for the automated detection and counting of *Aedes* eggs under real-world field conditions. The system operates directly on smartphone images without prior preprocessing, analyzes oviposition substrates up to 20.5 cm in length, and performs reliably in the presence of common field contaminants such as bio-attractant stains, sand, and particulate matter. Our system is accessible through a lightweight web interface and supported by a AI–based architecture that enables retraining for new ecological contexts or additional *Aedes* species. the system provides rapid and scalable egg quantification from smartphone images, strengthening ovitrap-based surveillance and community-level vector control.

## 2. Methodology

### 2.1. Biological sample collection

All analyzed samples (100%) consisted of eggs laid by wild *Ae. aegypti* females under natural field conditions. Eggs were collected on white nonwoven technical napkins (20.1 × 20.5 cm), which served as oviposition substrates. These napkins were placed lining the inner surface of handmade ovitraps with a capacity of 0.9 L. The traps were externally covered with black plastic film (vinyl wrap) to enhance visual contrast and stimulate oviposition, following previously described protocols [9,10]. The ovitrap prototype used in this study was developed by our team and is hereafter referred to as Ovicol[10]

To attract gravid females, a solution composed of 250 mL of tap water, five drops of a molasses solution (blackstrap molasses diluted 1:10 in tap water), and less than 0.5 g of dry yeast (*Saccharomyces cerevisiae*) was used. The yeast rapidly fermented the sugar, producing volatile compounds that act as attractants for ovipositing mosquitoes.

Ovicol were inspected every four days. During each cycle, 80 ovitraps were examined until a total of 400 samples was obtained. At every inspection, napkins were replaced and both the water and attractant solution were completely renewed.

In addition, 100 extra samples were collected using only tap water without bio-attractant. This modification aimed to reduce substrate staining caused by molasses fermentation and to evaluate system performance under conditions of lower visual contamination.

### 2.2. Reference manual counting and image acquisition

After collection, the napkins were processed fresh, and eggs were manually counted on a white plastic surface to obtain a preliminary human reference count under field conditions. This initial count served as an operational estimate during sample processing but was not used as the definitive ground truth for model training or evaluation.

Each sample was subsequently photographed using an iPhone X smartphone (12 MP real camera), capturing images at full resolution (3024 × 4032 pixels).

Images were then shared via WhatsApp messaging, which resulted in automatic compression and the introduction of JPEG artifacts. This procedure was intentional, as it realistically reproduces the operational conditions of fieldwork and community-level use, where images are commonly transmitted through mobile messaging platforms rather than preserved in raw format.

The final dataset consisted of 275 original images containing a total of 27,562 annotated eggs, with a mean of 99 eggs per image (range: 0–1052). Each image was labeled with the date of collection and the total number of eggs observed.

### 2.3. Annotation platform and virtual manual counting

To avoid reliance on external software and prevent loss of access to essential flat files required for AI model training (e.g., YOLO format in json), we developed a dedicated web-based platform for virtual manual annotation and counting of *Ae. aegypti* eggs, named Ovilab, available at https://ovilab.chilloa.org/.

The platform enables users to upload images, perform manual annotations directly on oviposition substrates, and store data centrally in the cloud. This architecture ensures traceability, quality control, and reusability of the dataset for model retraining and validation processes.

All definitive annotations used as ground truth for model training and evaluation were generated through the Ovilab platform using the digital images. This virtual annotation process allowed precise localization and counting of eggs while maintaining a structured and reproducible dataset.

All annotations were performed by a medical entomology expert to ensure consistency and accuracy in ground truth labeling. The software is open access, facilitating adaptation, transparency, and reuse for academic and scientific purposes (Fig 1).

**Fig 1.**
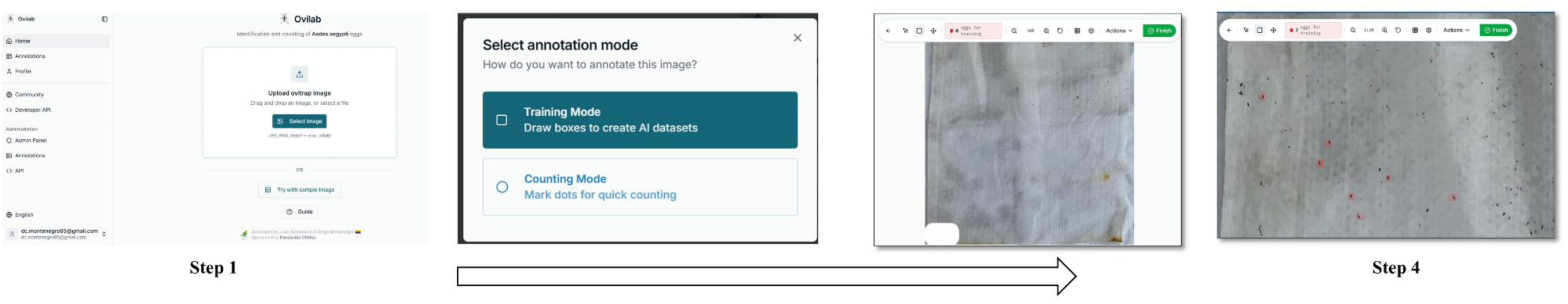
Ovilab web interface used for *Ae. aegypti* egg annotation and dataset generation for algorithm training

### 2.4. Image preprocessing and tiling strategy

Given the high resolution of the images (up to 3780 × 4032 pixels) and the small size of the eggs (≈10 pixels), a systematic tiling strategy was implemented. Each image was divided into 640 × 640 pixel tiles with 20% overlap between adjacent tiles. Tiles were created with a stride of 512 pixels (80% of tile size). This overlap ensured that eggs located near tile borders would appear fully visible in at least one cropped segment. For tiles corresponding to the edges of the original image, zero-padding (black borders) was applied to maintain uniform dimensions. Original annotations were transformed into the coordinate system of each tile, retaining only those instances in which at least 25% of the egg area remained visible within the cropped region. To prevent overfitting to background patterns, negative tiles (containing no annotations) were included at a 1:4 ratio relative to positive tiles, randomly sampled from available empty tiles for each image. This process generated a total of 2,753 tiles derived from the 275 original images. To ensure reproducibility and prevent data leakage, images were split at the source level (not tile level) using a fixed random seed (42), allocating 70% to training, 15% to validation, and 15% to testing subsets, ensuring tiles from the same original image never appeared across different splits.

### 2.5. Detection Model Training

A YOLOv11m model [17] was trained for 150 epochs using the Stochastic Gradient Descent (SGD) optimizer, with an initial learning rate of 0.01 and a batch size of 8. Early stopping was implemented with a patience of 30 epochs to prevent overfitting. A 5-epoch warmup phase was included to stabilize the initial stages of training.

To enhance robustness under highly variable real-world field conditions, an intensive data augmentation strategy was applied. This included full 360° rotation, 50% scale variation, horizontal and vertical flips (each with 50% probability), mosaic augmentation, mixup (α = 0.3), copy-paste augmentation (30% probability), random erasing (40% probability), and HSV color space adjustments (hue ±0.015; saturation ±0.7; value ±0.4). These transformations were designed to simulate common field-related artifacts such as lighting variation, surface contamination, irregular contrast, and arbitrary substrate orientation.

To improve localization performance for small objects—considering that eggs occupy approximately 10–15 pixels in the original images—the bounding box loss weight (box loss) was increased from 7.5 to 12.0. During inference, Non-Maximum Suppression (NMS) was applied using a confidence threshold of 0.15 and an Intersection over Union (IoU) threshold of 0.5, with a maximum of 300 detections allowed per image.

### 2.6. Inference and Count Reconstruction

During inference, complete images are divided into non-overlapping 640 × 640 pixel tiles for computational efficiency. Each tile is processed independently, and detections are mapped back to the original image coordinate system. Global Non-Maximum Suppression (IoU = 0.3) is applied to eliminate duplicate detections at tile boundaries, retaining only detections with a confidence level of 0.25 or higher.

### 2.7. Computational Infrastructure

Training was performed on an NVIDIA RTX 5090 GPU (32 GB VRAM) with an AMD EPYC 9654 96-core processor and 64 GB RAM running Ubuntu 24.04.3 LTS, completed in approximately 36 minutes for 150 epochs. The implementation was developed in Python v3.12.1[18] using the Ultralytics framework for YOLOv11m.

### 2.8. Statistical Comparison Between Egg Counting Methods

An independent subset of 30 samples, not used during Col-Ovo model training, was selected for comparative statistical validation. The Ovilab platform was defined as the reference standard, as it allows image magnification up to 200%, facilitating differentiation of fragmented and clustered eggs and thereby reducing the likelihood of underestimation or overestimation (S1 File).

Each sample was evaluated using three counting methods: i) Ovilab platform (reference standard), ii) Direct visual counting by a human expert, and iii) The Col-Ovo AI system.

Additionally, the expert classified each sample into three levels of difficulty (1–3), with level 3 corresponding to the highest degree of contamination (e.g., dust, sand, debris, or artifacts potentially confusable with *Ae. aegypti* eggs).

Because repeated measurements were obtained from the same samples and normality of counts was not assumed, non-parametric tests were applied: i) Global comparison among methods (Friedman test), ii) Stratified comparison by difficulty level (Friedman test by subgroup), iii) Pairwise comparisons (Wilcoxon signed-rank test), iv) Association between methods (Spearman correlation), and v) Agreement analysis (Bland–Altman method).

To further evaluate interchangeability between methods, Lin’s concordance correlation coefficient (ρc) was calculated, integrating both precision (correlation) and accuracy (closeness to the line of identity). This analysis was performed globally and stratified by difficulty level to assess potential performance degradation under increasingly complex conditions.

Effect sizes were calculated as Kendall’s W for the Friedman test and Wilcoxon’s r (Z/√n) for pairwise comparisons. Statistical significance was defined as p < 0.05.

Finally, processing time per sample was compared between Col-Ovo and the Ovilab platform by measuring differences in seconds and estimating the relative increase in operational throughput.

All statistical analyses were conducted using open-source Python v3.12.9[18], within the interactive Jupyter Notebook v7.3.2 environment [19] for data processing, modeling, and evaluation.

## Results

### 1. Performance of the Col-Ovo Model

To evaluate the effectiveness of the proposed automated egg-counting approach, we assessed the performance of the Col-Ovo model. The Col-Ovo for *Ae. aegypti* eggs achieved a mAP@0.5 of 0.825 and a mAP@0.5:0.95 of 0.332, with a precision of 0.804, recall of 0.794, and an F1-score of 0.80. These metrics indicate a balanced performance between accurate detection and completeness, maintaining a low rate of false positives while preserving a robust ability to recover eggs present in the images.

### 2. Efficiency of Col-Ovo Compared with the Ovilab Standard and Human Counting

In the subset of 30 independent samples, the Friedman test revealed marginally significant differences among counting methods (χ² = 5.81; p = 0.0548), suggesting that at least one pair of methods differed slightly. However, the overall level of discordance was low (Kendall’s W = 0.097), indicating limited global disagreement across methods.

When counts were stratified by difficulty level, no statistically significant differences were observed between methods (all p > 0.07), suggesting consistent performance across increasing levels of sample complexity.

Pairwise comparisons using Wilcoxon signed-rank tests showed that Human counting differed significantly from the Ovilab reference standard (*p* = 0.009; effect size *r* = 0.48), while Col-Ovo demonstrated comparable performance to both the reference standard (*p* = 0.252; *r* = 0.21) and Human counting (*p* = 0.273; *r* = 0.20). These findings indicate that Col-Ovo is statistically indistinguishable from the reference method, whereas Human counting exhibits systematic deviation (Fig 2).

**Fig 2.**
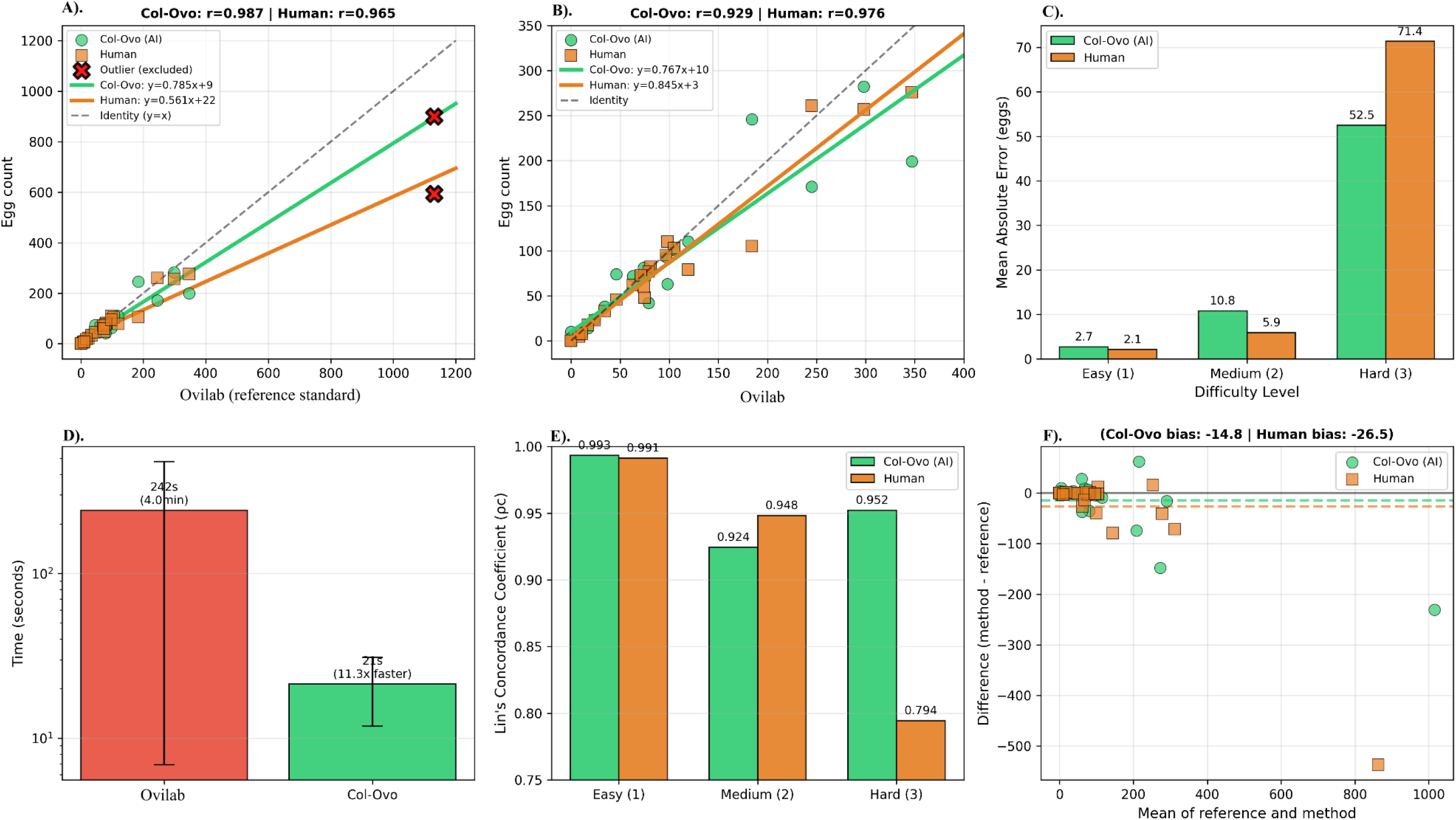
Comparative performance of Col-Ovo (AI) and Human counting against the Ovilab reference standard. A) Regression with outlier (n=30): Col-Ovo (green) shows higher correlation and steeper slope (0.785) than Human (orange, slope 0.561). Dashed line = identity (y=x). (B) Regression without outlier (n=29, 0–400 eggs range). (C) Mean absolute error by difficulty level. (D) Processing time comparison (log scale). (E) Lin’s concordance coefficient by difficulty. (F) Bland-Altman plots: Col-Ovo (green circles) shows narrower limits of agreement and lower bias than Human (orange squares).

Linear regression analysis using Ovilab as the independent variable revealed differential performance between methods. Col-Ovo showed a slope of 0.785 (95% CI: 0.72–0.85) with intercept 8.9, explaining 97.4% of variance (*R²* = 0.974). Human counting exhibited a shallower slope of 0.561 (95% CI: 0.48–0.64) with intercept 21.7, explaining 93.1% of variance (*R²* = 0.931). The deviation from unity slope (ideal: 1.0) indicates that both methods underestimate high egg counts, but Col-Ovo maintains better proportionality (21.5% vs. 43.9% underestimation) (Fig 2A and Fig 2B).

Bland-Altman analysis confirmed higher agreement for Col-Ovo relative to Human counting (Fig 2F). Col-Ovo exhibited narrower limits of agreement (LoA₉₅%: −119.1 to +89.5 eggs) compared to Human counting (LoA₉₅%: −220.2 to +167.2 eggs), with lower systematic bias (mean difference: −14.8 vs. −26.5 eggs) and lower mean absolute error (23.4 vs. 28.6 eggs).

Stratified analysis by difficulty level revealed Col-Ovo’s superior robustness under challenging conditions. Lin’s concordance coefficient (ρc) remained high across all levels: Easy (ρc = 0.993), Medium (ρc = 0.924), and Hard (ρc = 0.952) (Fig 3). In contrast, Human counting showed marked degradation at the Hard level (ρc = 0.794), representing a 16% decline in concordance compared to Col-Ovo (Fig 2E). Mean absolute error in Hard samples was 26% lower for Col-Ovo (52.5 vs. 71.4 eggs).

**Fig 3.**
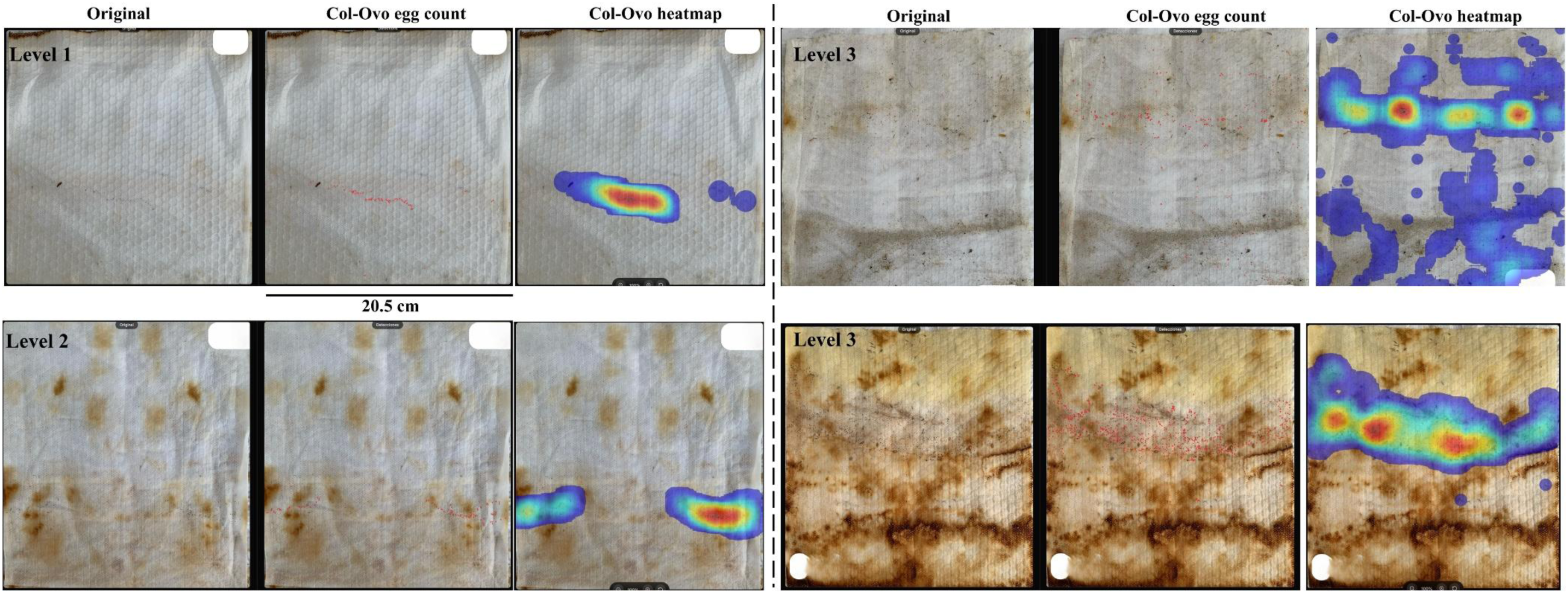
Stratification of oviposition substrates (20.5 cm) according to human difficulty in egg recognition and counting: Level 1 (Easy), Level 2 (Medium), and Level 3 (Hard), and their corresponding processing by Col-Ovo.

Col-Ovo demonstrated substantial time efficiency, processing samples in 21 seconds (SD = 9.5) versus 242 seconds (SD = 236; 4.0 minutes) for the reference Ovilab method—an 11.3-fold speed increase (throughput: 3.2 vs. 0.25 samples/minute). This efficiency advantage was maintained across all difficulty levels without accuracy degradation.

### 3. Col-Ovo Availability and Interface

Col-Ovo is freely accessible as a web-based application at https://colovo.chilloa.org/en/, requiring no software installation or specialized hardware beyond a standard internet-connected device. The platform implements a five-step streamlined workflow designed for field operability (Fig 4):

**Fig 4.**
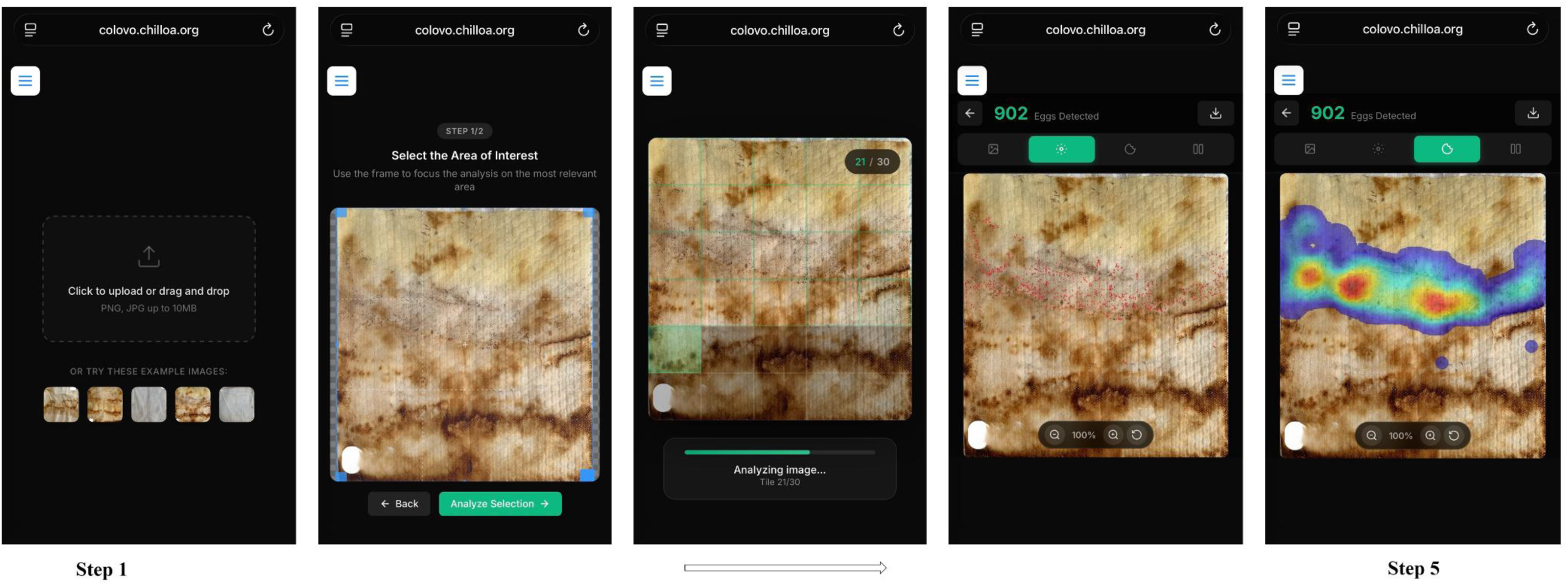
Col-Ovo web interface and step-by-step workflow for automated counting of *Ae. aegypti* eggs.

Step 1: Image Upload. The interface presents a drag-and-drop upload area supporting PNG and JPG formats up to 10MB. Pre-loaded example images are available for users to test system functionality without requiring their own samples.

Step 2: Region of Interest Selection. Users can adjust a resizable frame to focus analysis on the most relevant substrate area, eliminating background noise and optimizing computational resources. This feature is particularly valuable for heterogeneous substrates up to 22 cm in length.

Step 3: Automated Analysis. The system processes the image through a grid-based tile analysis, with an average of 30 tiles per image (as shown in the progress indicator), providing real-time status updates (e.g., “Analyzing image… Tile 30/30”). Processing completes in 21 seconds per image on average (SD = 9.5)

Step 4: Results Visualization. The interface displays the total egg count (e.g., “902 Eggs Detected”) with multiple visualization modes:

- Detection view: Individual egg locations marked with red dot annotations
- Heatmap view: Density visualization using color gradients (blue to red) indicating high-probability egg clusters
- Zoom and pan controls (100% default magnification with adjustable scale)

Step 5: Data Export. Results can be downloaded for record-keeping and integration with surveillance databases.

The interface prioritizes usability for non-specialist personnel, eliminating the need for extensive training in entomological identification. The system processes samples at 3.2 samples/minute, representing an 11.3-fold throughput increase compared to manual reference methods (0.25 samples/minute). Importantly, accuracy is maintained across all difficulty levels, as evidenced by high concordance even in hard samples with contamination (Lin’s ρc = 0.952).

## Discussion

Here we show that Col-Ovo enables the accurate, rapid, and scalable automated counting of *Ae. aegypti* eggs from smartphone images captured under routine field conditions. This capability addresses a critical operational bottleneck in ovitrap-based surveillance. The egg stage represents a particularly strategic intervention point in the biology of *Aedes* mosquitoes: eggs can withstand desiccation for several months, facilitate long-distance dispersal through human transport[6,20], and can be effectively targeted through community-based strategies using bioattractants and inexpensive oviposition substrates. Enabling rapid egg quantification therefore strengthens the practical implementation of egg-focused vector control strategies. Our findings indicate that Col-Ovo achieves accuracy comparable to the reference annotation platform (Ovilab) while reducing the variability associated with manual counting. Overall differences among methods were minimal (Friedman χ² = 5.81; p = 0.0548; Kendall’s W = 0.097), but pairwise comparisons showed that manual counting differed significantly from the reference platform (p = 0.009), whereas Col-Ovo showed no significant differences relative to either the reference method (p = 0.252) or manual counting (p = 0.273). These results suggest that automated detection can reproduce the reliability of standardized digital annotation while minimizing observer-dependent variability.

Performance differences became more evident under conditions of high egg density. Although both manual and automated approaches tended to underestimate counts as egg aggregation increased, similar limitations have been reported for previous automated tools [21]. However, Col-Ovo maintained substantially better proportional agreement with the reference standard (slope = 0.785; R² = 0.974) compared with manual counting (slope = 0.561; R² = 0.931), reducing proportional underestimation by approximately 50%. Stratified analyses further supported this interpretation: concordance remained stable for Col-Ovo across difficulty levels (Lin’s ρc > 0.92), whereas manual counting deteriorated in high-density samples (ρc = 0.794). These findings indicate that automated detection is less susceptible to aggregation-related counting fatigue, a known limitation of manual ovitrap surveillance.

A major advantage of Col-Ovo lies in its operational accessibility. The system can reliably analyze images captured with standard smartphones, including those compressed through common messaging platforms such as WhatsApp. It is publicly available, compatible with widely used mobile devices, and independent of specialized hardware, allowing egg counting to be performed directly from images obtained under routine field conditions. This design substantially reduces the technological barriers that often limit the implementation of AI-based tools in endemic regions.

Integration with the OviLab annotation platform further strengthens this framework by enabling structured annotation, centralized storage of labeled datasets, and model retraining using additional image repositories [22]. This architecture improves data traceability and reproducibility while creating opportunities to expand automated detection toward taxonomic differentiation at the egg stage, as suggested by recent morphometric and microscopic studies [23,24].

Operational efficiency represents an additional advantage. Col-Ovo processed samples in approximately 21 seconds compared with 242 seconds required for the OviLab workflow, representing an 11.3-fold increase in throughput while maintaining statistical concordance. Such improvements may substantially increase the analytical capacity of vector surveillance programs, particularly in settings where large ovitrap networks must be processed with limited personnel.

These findings should also be interpreted in light of the technical challenges inherent to field-based detection of mosquito eggs. *Ae. aegypti* eggs are extremely small (∼0.065 mm²) and frequently occur on heterogeneous substrates stained by bioattractants and photographed under uncontrolled lighting conditions[25]. Under these conditions, the difference between mAP@0.5 (0.825) and mAP@0.5:0.95 (0.332) reflects expected trade-offs in detecting clustered eggs (Gaburro et al., 2016; S1 Fig), where partial overlap and variable orientation reduce bounding-box agreement under strict IoU thresholds.

Previous attempts to automate egg detection illustrate these challenges. Early segmentation-based approaches achieved relatively low counting error but relied on extremely small training datasets obtained under laboratory conditions [14,26]. Subsequent deep learning systems improved detection performance but often introduced operational constraints. For example, ICount showed reduced accuracy in clustered egg scenarios[21], whereas hardware-assisted systems required specialized scanners or imaging equipment [16]. More recent models such as EggCountAI reported high correlation with manual counts but relied primarily on laboratory images and computer-based processing[12].

In contrast, Col-Ovo was designed specifically for field conditions. By operating directly on smartphone images and eliminating the need for specialized hardware, the system substantially lowers barriers to implementation in endemic regions. Combined with the OviLab annotation framework, this approach enables continuous model improvement and facilitates the integration of automated egg detection into routine surveillance workflows.

From a public health perspective, the ability to automate egg quantification has important implications for vector surveillance and control. Ovitrap networks are widely used as early indicators of *Aedes* activity [8], but their operational value is frequently constrained by the time and training required for manual counting. By reducing analysis time from minutes to seconds, Col-Ovo enables faster interpretation of ovitrap data and facilitates monitoring of large surveillance networks. This acceleration may improve outbreak preparedness by enabling earlier detection of increases in vector density and more timely identification of transmission hotspots.

More broadly, the integration of low-cost oviposition substrates, community participation, and automated egg detection may strengthen preventive approaches to vector control. In the context of the continued global expansion of *Aedes* mosquitoes and the increasing burden of arboviral diseases, accessible and hardware-independent AI tools such as Col-Ovo represent an important step toward scalable surveillance systems capable of supporting community-centered vector management strategies.

---

**S1 Fig.**
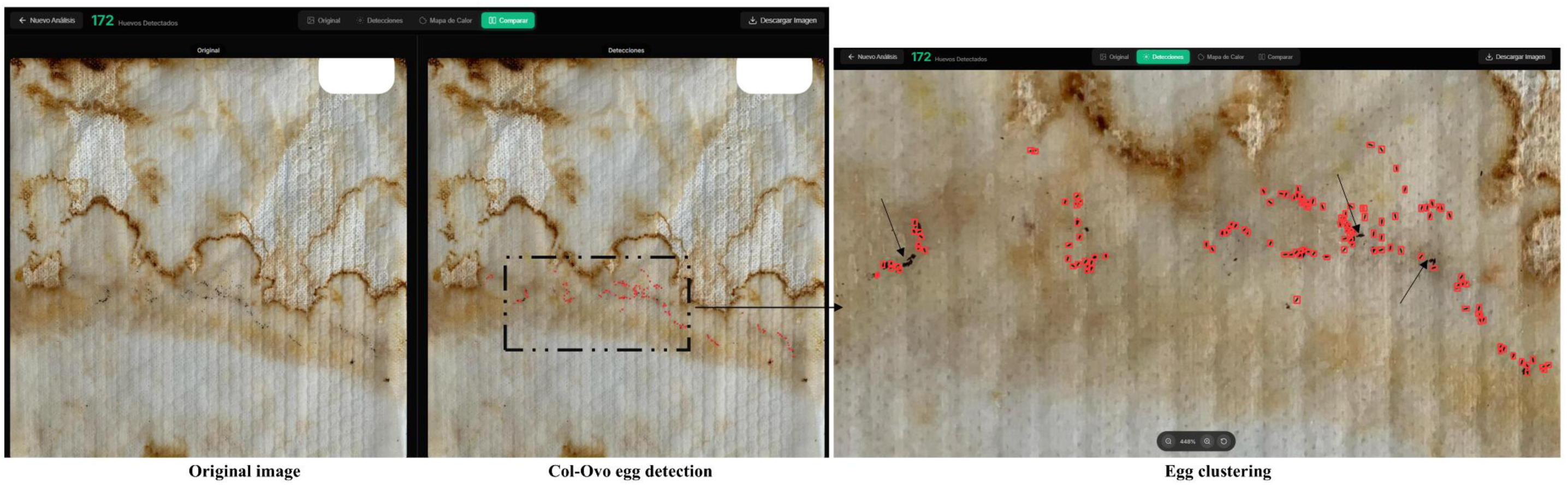
Oviposition substrate with clustered *Ae. aegypti* eggs representing a challenging scenario for automated detection by the Col-Ovo AI system.

## Acknowledgments

We thank the technical team of the Programa Regular de Vigilancia y Control de Enfermedades Transmitidas por Vectores (Vector-Borne Diseases Surveillance and Control Program) of Santa Marta for their support in field activities. We also thank Manel Camps and Andres Rico for their conceptual review of the manuscript.

## Author contributions

JA was responsible for the development of the OviLab and Col-Ovo platforms. DM designed and conceptualized the study, conducted field data collection, trained the Col-Ovo model, performed the analyses, and led the manuscript writing. Both authors contributed to manuscript review and approved the final version.

## Ethics approval and consent to participate

Not applicable, as this work was conducted within the legal mandates and public health competencies of the respective health authorities (Vector-Borne Diseases Surveillance and Control Program of Santa Marta).

## Consent for publication

Not applicable.

